# Low-CO_2_ inducible bestrophins in diatom thylakoid membranes sustain high photosynthetic efficacy at distant locations from the pyrenoid

**DOI:** 10.1101/2023.09.20.558591

**Authors:** Minori Nigishi, Ginga Shimakawa, Kansei Yamagishi, Ryosuke Amano, Shun Ito, Yoshinori Tsuji, Chikako Nagasato, Yusuke Matsuda

## Abstract

Anion transporters are important to sustain a variety of physiological states in cells. Bestrophins are a family of Cl^−^ and/or HCO3^−^ transporters conserved in bacteria, animals, algae, and plants. Recently, bestrophin paralogs were found in the green alga *Chlamydomonas reinhardtii* as up- regulated components in low CO_2_ conditions that play an essential role in the CO_2_- concentrating mechanism (CCM). Bestrophin orthologs are also conserved in diatoms, a group of secondary endosymbiotic algae harboring red-type plastids, but their physiological functions are not known yet. Here, we characterized the subcellular localization and expression profile of bestrophins in the marine diatoms *Phaeodactylum tricornutum* (PtBST1−4) and *Thalassiosira pseudonana* (TpBST1 and 2). PtBST1 and PtBST2 were localized at the stromal thylakoid membrane outside of the pyrenoid, and PtBST3 was localized in the pyrenoid. Contrarily, TpBST1 and TpBST2 were both localized in the pyrenoid. These bestrophin proteins were accumulated in cells grown in atmospheric CO_2_ but not in 1% CO_2_-grown cells. To assess the physiological functions, we generated knock-out mutants for PtBST1 by genome editing. The lack of PtBST1 decreased affinity of photosynthesis for dissolved inorganic carbon closer to that of the cells grown in 1% CO_2_. Additionally, non-photochemical quenching was 1.5–2.0 times higher in the mutants than that of the wild type cells. These data suggests that HCO3^−^ transport at the stroma thylakoid membranes by PtBST1 is a critical part of the CO_2_ evolving system of the pyrenoid in the fully induced CCM, and simultaneously that PtBST1 modulates photoprotection in response to CO_2_ availability in *P. tricornutum*.

**Significant statement:** Marine diatoms are responsible for nearly half of oceanic primary production, owing to the high-affinity photosynthesis for dissolved inorganic carbon which is supported by CO_2_- concentrating mechanism (CCM). This study uncovered that a bestrophin family protein at the stoma thylakoid membrane operates to import HCO_3_^−^ to the thylakoid lumen and mobilizes it towards the CO_2_ evolving system at the pyrenoid-penetrating thylakoid in the diatom *Phaeodactylum tricornutum*. This HCO_3_^−^ collecting system not only enhances the CCM but also down regulates the photoprotection capacity of photosystem II, presumably by affecting the thylakoid lumen acidification. This study experimentally demonstrates the molecular mechanism how diatoms optimize the use of CO_2_ and light energy, giving an insight into the reason of ecological successfulness of marine diatoms.

## Introduction

Marine diatoms are a diverse class of marine phytoplankton that are widely spread in the global ocean and responsible for nearly half of oceanic primary production (Falkowski et al., 1998). The diatom lineage arose due to a secondary endosymbiotic event, with diatom cells harboring chloroplasts originally derived from red algae. As is typical in secondary endosymbionts, the diatom chloroplast is comprised of four-layered membranes; two outer layers termed the chloroplast endoplasmic reticulum (CER) followed by two layers termed the chloroplast envelope (CE). Between the CER and CE, there is a remnant space left over from the cytosol of the ancestral red algal called the periplastidal compartment (Flori et al., 2016). Within the diatom stroma there is a triplet layered thylakoid membrane called the girdle lamella with the layered stromal thylakoid (ST) at its interior. In the middle of the stroma, there is a phase separated proteinaceous body called the pyrenoid (Bedoshvili et al., 2009).

To sustain high efficacy CO_2_ fixation under sea water conditions where the CO_2_ concentration is low (∼10 µM) but HCO_3−_ is predominant (> 2 mM), diatoms actively take up external HCO_3−_ and/or facilitate diffusive entry of CO_2_ dehydrated from external HCO_3−_ by carbonic anhydrases (CAs) expressed in the periplasm (Nakajima et al., 2013; Nawaly et al., 2022). Dissolved inorganic carbon (DIC) taken up from sea water is presumably further mobilized into the chloroplast by an active HCO_3−_ transport system on the chloroplast membranes (Matsuda et al., 2017). Finally, DIC in the stroma is converted into CO_2_ at the pyrenoid where the ribulose 1,5-bisphosphate carboxylase/oxygenase (Rubisco) enzyme is condensed (Tsuji et al., 2017). This overall system is termed the biophysical CO_2_-concentrating mechanism (CCM).

The pyrenoid is surrounded by layers of ST and the outermost girdle lamella where the photosynthetic electron transport system produces NADPH and ATP to drive the Calvin cycle. A part of thylakoid membranes traverses through the pyrenoid matrix, which is denoted here as the pyrenoid-penetrating thylakoid (PPT) membrane. In the pennate diatom *Phaeodactylum tricornutum*, the θ-type carbonic anhydrase, Ptθ-CA1, occurs at the PPT lumen where light-driven-acidification accelerates the dehydration of HCO_3−_ to evolve CO_2_ (Kikutani et al., 2016). Similar machinery appears to occur also in the centric diatom *Thalassiosira pseudonana*, because the PPT luminal θ-type CA was recently identified (Nawaly et al., 2023). The CA-containing PPT at the pyrenoid core appears to be the fundamental structure of the CO_2_- evolving machinery conserved in marine diatoms, whereas the strategies to acquire HCO_3−_ from sea water are likely diverse among diatoms. One of the critical missing links to understand the diatom CCM is identifying the system to facilitate influx of HCO_3−_ across the thylakoid membranes and into the PPT lumen.

The CO_2_-evolving machinery coupled with luminal CA is also conserved in the pyrenoid of the green alga *Chlamydomonas reinhardtii*, strongly suggesting the occurrence of convergent evolution across distant taxa of algae. Given the steady and continual decrease in atmospheric CO_2_ concentration over Earth’s history, such a system would have been highly selected for to optimize the use of light and CO_2_ in the aquatic environment. It is recently reported that the bestrophin paralogs BST1−3 localized in the thylakoid membranes transport HCO_3−_ to support the CO_2_-evolving reaction by the α-type CA, CAH3, in the lumen of the pyrenoid-invaginating thylakoid membranes (termed as the pyrenoid tubule) in *C. reinhardtii* (Mukherjee et al., 2019), most probably as a part of the CO_2_-evolving machinery in *C. reinhardtii*. Transcripts of BST1−3 in *C. reinhardtii* were highly accumulated in the cells grown in atmospheric CO_2_, but were found only in trace amount in 5% CO_2_-grown cells, clearly indicating the low-CO_2_-inducible nature of these BSTs (Mukherjee et al., 2019). The BSTs expression in *C. reinhardtii* probably occurs in concert with the structural dynamics of the pyrenoid in response to CO_2_ that is required to play a specific role in the CCM under CO_2_ limitation. Even though the function of these BSTs seemed tightly related to the function of the pyrenoid tubules, the location of these BSTs in low-CO_2_-grown *C. reinhardtii* was not likely limited to the pyrenoid areas but also localized in the thylakoid membranes other than the pyrenoid tubule (Mukherjee et al., 2019).

Bicarbonate is an anion and thus could counteract the proton-derived thylakoid membrane potential. Conversely, a high activity of HCO_3−_ dehydration catalyzed by luminal CAH3 in the pyrenoid tubules results in consumption of protons, potentially dissipating the proton gradient across the membranes (ΔpH). These potential effects of the introduction of HCO_3−_ into the thylakoid lumen could impose a significant impact on the function of the photosynthetic electron transport. The problem of the effect of HCO_3−_ introduction into the lumen of the spatially and functionally diverse thylakoid membrane system is still an open question in *C. reinhardtii*. Diatoms also possess putative bestrophin sequences in their genome, but the function of these putative candidates, localizations, or expressional controls is not yet totally known.

Here, we searched putative bestrophin genes and found four and two isoforms in *P. tricornutum* and *T. pseudonana*, respectively. These putative bestrophins were localized in the different chloroplast areas and highly expressed in response to CO_2_ limitation in regulation at either transcriptional or post-transcriptional processes. The most abundant isoform in *P. tricornutum*, denoted as PtBST1, was targeted by a highly specific genome editing (CRISPR/Cas9 nickase) to generate the knock-out mutants. The photosynthetic phenotypes of the mutants suggest that PtBST1 functions as the HCO_3−_ transporter in the low-CO_2_ inducible CCM in *P. tricornutum*, and further modulates photoprotective mechanisms in relation to the utilization of H^+^ in the thylakoid lumen.

## Results

### Screening of BST orthologs in marine diatoms

Bestrophin is often defined as a Ca^2+^-activated anion transporter (Yang et al., 2014) and its sequences were clustered to several large protein families comprising a variety of organisms, including bacteria, animals, and plants. Indeed, a BLAST search for Chlamydomonas BST1−3 sequences found multiple bestrophin homologs in each species in diverse photosynthetic organisms, including cyanobacteria, green algae, red algae, land plants, and secondary algae (Supplementary Fig. S1). Among these homologs, the *C. reinhardtii* BST1−3 (Mukherjee et al., 2019) form a large conserved clade including land plants, cryptomonads, haptophytes, and diatoms. In this study, we defined the orthologs categorized into this clade as “BST” for the candidates of HCO_3−_ transporter in the thylakoid membranes.

BST orthologs are widely found in green algae and land plants, and secondary algae such as haptophytes and diatoms (Fig. 1A). Diatom BSTs further constituted a specific subclade that is apart from the subclade of the green linage. Meanwhile, cyanobacteria, red algae, and some secondary algae do not possess the BST orthologs. The BST homologs could not be recognized in the prasinophyceae algae *Ostreococcus*, a basal green algal lineage (Supplementary Fig. S1), suggesting that the BST in green lineage might have been acquired in the common ancestor of streptophytes and chlorophytes, and thereafter inherited to plants. It is likely that BST in secondary algae, such as cryptomonads, haptophytes, and diatoms, might be derived *via* a past temporal endosymbiotic association (cryptic endosymbiosis) with a green alga prior to the acquisition of present red-type chloroplast (Moustafa et al., 2009), whereas the BST homologs had been lost in a part of secondary algae. More than two BST isoforms were identified in seven diatom species whose genome information is presently available (Fig. 1B): pennate diatoms encode more than three BSTs, while centrics harbor two BST isoforms. In *P. tricornutum*, PtBST1 (Phatr3_J46336) and PtBST2 (Phatr3_J26635) share high similarity (approximately 66%) and formed a pennate-specific clade, whereas PtBST3 (Phatr3_J46366) and PtBST4 (Phatr3_J46360) were in separate group (Fig. 1B). *T. pseudonana* has two bestrophin paralogs (TpBST1, THAPSDRAFT_4819; and TpBST2, THAPSDRAFT_4820), which show approximately 66% similarity and are categorized into the centric-specific clade adjacent to the pennate-specific group that includes PtBST1 and 2 (Fig. 1B).

**Figure 1.**
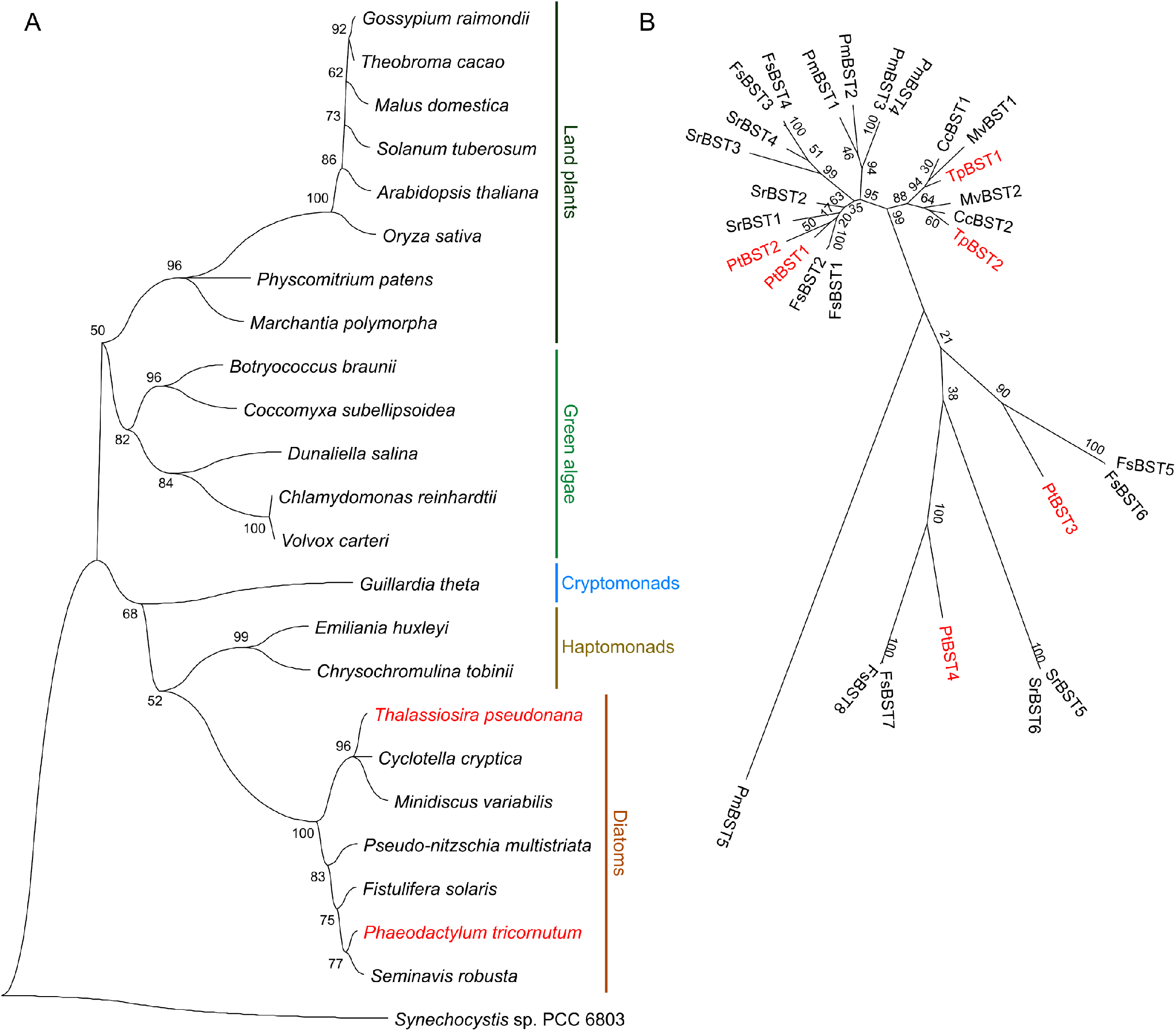
Phylogeny of BST isoforms in photosynthetic organisms. (A) Evolutionary relationship of BST orthologs in land plants, green algae, cryptomonads, haptophytes, and diatoms. The cyanobacterium *Synechocystis* sp. PCC 6803 is an outgroup. Bestrophin family genes are widely found in photosynthetic organisms, and one clade was analyzed as BST orthologs. One BST ortholog per species were selected for the phylogenetic analysis, based on the similarity to PtBST1. (B) Similarity of BST isoforms in seven diatom species. Species names were abbreviated as follows: *Phaeodactylum tricornutum* (Pt), *Thalassiosira pseudonana* (Tp), *Pseudo-nitzschia multistriata* (Pm), *Fistulifera solaris* (Fs), *Seminavis robusta* (Sr), *Cyclotella cryptica* (Cc), and *Minidiscus variabilis* (Mv).

One more BST isoform (Phatr3_J46365) was suggested by the database to occur in *P. tricornutum* showing 73% similarity to PtBST3, which could be denoted as PtBST5. Since the coding sequence is possibly derived from a splice variant or other allele to PtBST3, PtBST5 was excluded of the main subject in this study. Nevertheless, we note that the 3ʹ-extension region of PtBST3 is specifically lost in PtBST5, although these two BST isoforms show almost the same coding sequences (Supplementary Fig. S2).

### Subcellular localization of diatom BST isoforms

Based on the coding sequences from available databases, all these BST isoforms except for PtBST4 and TpBST1 were predicted to have the chloroplast transit peptide (Gruber et al., 2007). However, in this study, a 5ʹ-RACE analysis of PtBST4 and TpBST1 identified an upstream start codon that was missing in the database, and the chloroplast transit peptide was predicted in the newly obtained sequences of PtBST4 and TpBST1. Overall, all BST isoforms in *P. tricornutum* and *T. pseudonana* were predicted to be localized in chloroplasts (Supplementary Table S1).

To determine the specific localization in chloroplasts, we generated the transformants that express GFP-fused BST isoforms in *P. tricornutum* and *T. pseudonana*. The green fluorescence derived from PtBST1:GFP and PtBST2:GFP was detected across the whole chloroplast (Fig. 2A). The localization of PtBST1:GFP was further analyzed by electron microscopy with a GFP specific antibody (Fig. 2B), clearly showing its localization within the thylakoid membrane outside of the pyrenoid. These data suggest that PtBST1 and PtBST2 are localized mainly in the ST membrane outside of the pyrenoid. Meanwhile, the green fluorescence derived from GFP-fused PtBST3 was recognized specifically in a center part of chloroplasts where chlorophyll fluorescence was absent (Fig. 2A), an area most probably of the pyrenoid, strongly suggesting that PtBST3 occurs in PPT. To reveal a sequence motif that localizes PtBST3 to the PPT, we further analyzed the subcellular localization of a truncated PtBST3 lacking in the C-terminal extension sequence (G407−P519) that specifically occurs in PtBST3 but not in PtBST5 (Fig. S2) by expressing as the GFP fusion protein. The truncated PtBST3^Δ407-519^:GFP altered its location and the GFP signal was detected throughout the chloroplast in contrast to PtBST3:GFP, suggesting that the C-terminal region of PtBST3 contains the key factor targeting the protein to the PPT in *P. tricornutum* (Fig. 2A). Unfortunately, we could not obtain the transformant that expresses PtBST4:GFP in this study.

**Figure 2.**
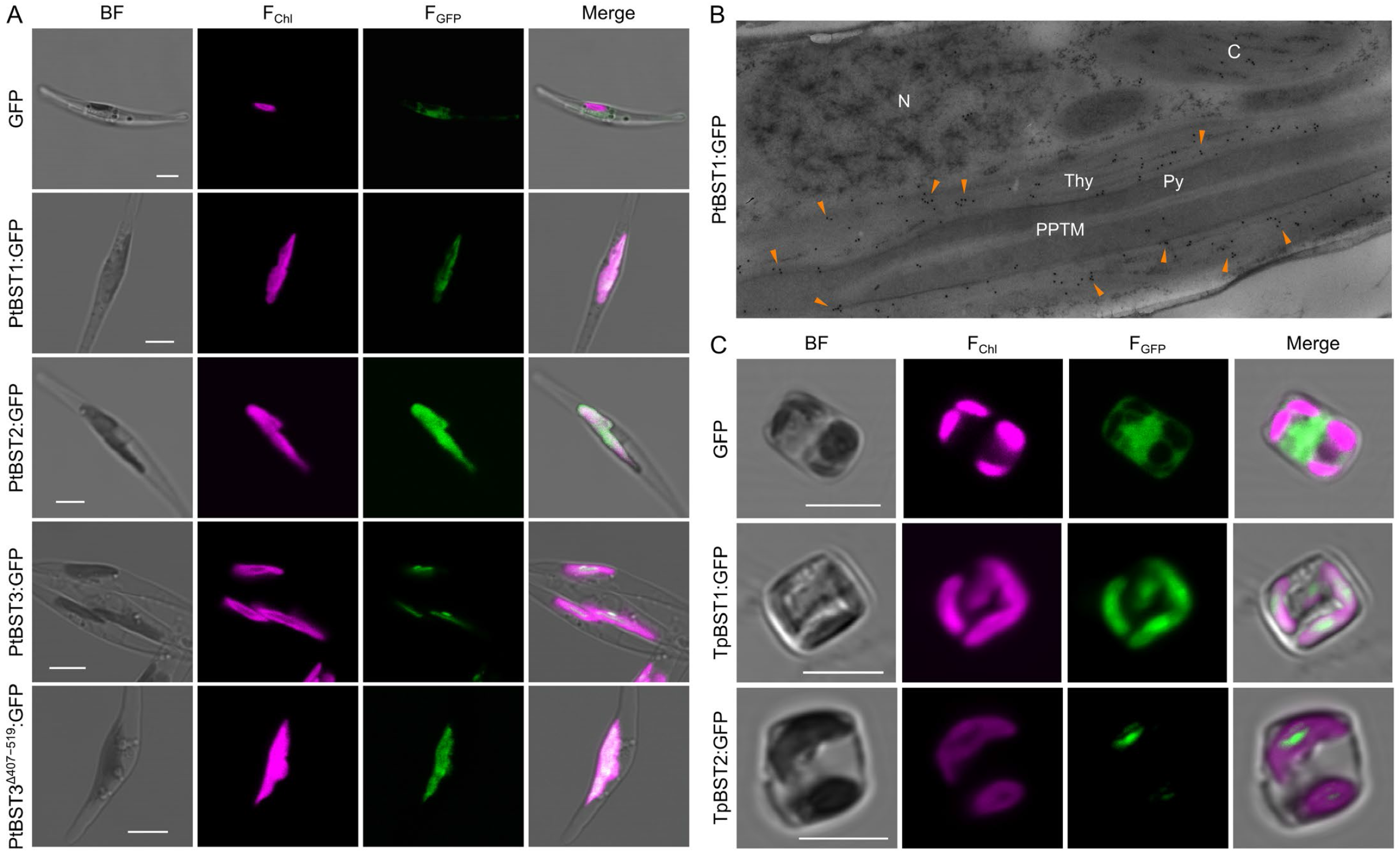
Subcellular localization of BST isoforms in *P. tricornutum* (A, B) and *T. pseudonana* (C). (A, C) Confocal images of bright field (BF), chlorophyll autofluorescence (F_Chl_), green fluorescence derived from GFP tagged to BST (F_GFP_), and all merged. White bars indicate 5 µm. (B) Immunoelectron microscopy with a specific antibody to GFP in the *P. tricornutum* transformant that expresses GFP-tagged PtBST1. N, nuclei; C, chloroplasts; Thy, stromal thylakoid membranes; Py, pyrenoid; PPT, pyrenoid-penetrating thylakoid membranes.

In *T. pseudonana*, both TpBST1 and TpBST2 were found to be localized in the PPT or around the pyrenoid according to the GFP signal of respective GFP fusion proteins (Fig. 2C), which is consistent with the latest data of TpBST2 localization by Nam et al. (Nam et al., 2022). Compared with TpBST2, the green fluorescence of TpBST1:GFP was detected broadly in the chloroplast (Fig. 2C). TpBST1 thus might be localized in both ST and PPT.

### Responses of the expression of diatom BSTs to changing CO_2_ concentrations

It has been reported that the CCM components are highly expressed under CO_2_ limitation in most cases at transcript levels in marine diatoms (Ohno et al., 2011; Nakajima et al., 2013; Hennon et al., 2015). Here, we found three “TGACGT/C” motifs, which is a target of a basic zipper (bZIP) 11 transcription factor, in the upstream genome sequence of *PtBST1* (Supplementary Fig. S3). The similar motif set has been previously characterized in the promoter region for the low-CO_2_-inducible β-CA1 gene, *PtCA1* in *P. tricornutum* and defined as the CO_2_-cAMP-responsive elements (Ohno et al., 2011). Therefore, PtBST1 was expected to be derepressed at high CO_2_ concentration *via* the cAMP-dependent signalling pathway. Additionally, in *T. pseudonana*, both TpBST1 and TpBST2 have been found to be up-regulated under CO_2_ limitation in a proteomic analysis (Kustka et al., 2014).

Here, we examined the expression levels of four and two BST isoforms in *P. tricornutum* and *T. pseudonana*, respectively. Compared with 1% CO_2_ (high CO_2_, HC) condition, cells grown under an air-level CO_2_ (LC) condition highly accumulated *PtBST1−4* transcripts (Fig. 3A). In particular, the transcript level of *PtBST1* was significantly higher (10- 20 times) than those of other *PtBSTs* and the induction ratio of *PtBST1* transcript in LC over HC was more than 40 times (Fig. 3A), strongly suggesting that PtBST1 is the most abundant BST isoform in *P. tricornutum*. The accumulation of PtBST1 at the protein level was also investigated by immunoblotting with newly generated anti-PtBST1 antiserum. Consistent with the transcript level, PtBST1 protein was highly expressed in the LC condition, while in HC-grown cell lysate, PtBST1 was found only in trace amounts (Fig. 3B). In *T. pseudonana*, in sharp contrast to *P. tricornutum*, the abundance of the *TpBST1* and *TpBST2* transcripts was not different between HC and LC-grown cells, and the transcript of *TpBST2* was even lower in the LC condition (Fig. 3C), indicating that the expression of TpBSTs was not controlled at the transcript level. However, interestingly, both TpBSTs isoforms were never detected at the protein level in the cells grown in HC but highly accumulated in LC-grown cells (Fig. 3D). These data strongly suggest that the expressions of BSTs in *T. pseudonana* are induced by CO_2_ limitation at the post transcriptional levels; in other words, translation of these *TpBSTs* transcripts is almost completely suppressed in HC. Given these data, the regulatory mechanisms of BST expression in response to CO_2_ concentrations between these two diatom species are totally different.

**Figure 3.**
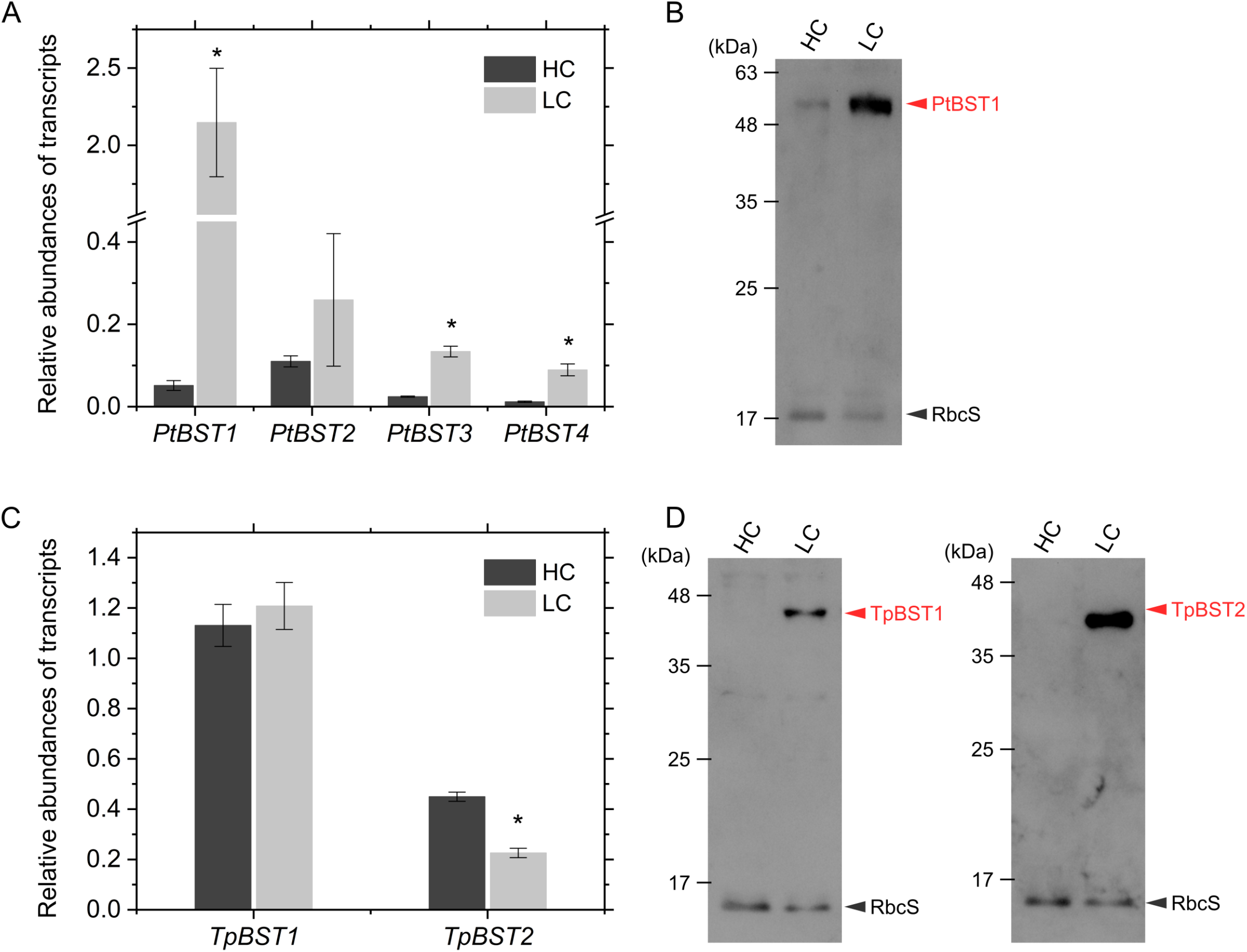
Expressions of BST isoforms under air-level (LC) and 1% (HC) CO_2_ in *P. tricornutum* (A, B) and *T. pseudonana* (C, D). (A, C) Relative abundances of transcripts estimated by qRT- PCR with *Actin1* as the reference gene. Data are shown as the mean ± standard deviation (*n* = 3, biological replicates). Asterisks indicate statistically significant differences (*p* < 0.05) between HC and LC as per Student’s *t*-test. (B, D) Accumulation of PtBST1, TpBST1, and TpBST2 proteins estimated by immunoblotting with each specific antibody. Whole crude extracts (10 µg protein each) were analyzed. RbcS protein was used as the loading control. Representative data of three independent experiments are shown.

### Highly specific genome editing to PtBST1 using CRISPR/Cas9 nickase

In this study, we performed the gene disruptions of *PtBST1* in *P. tricornutum* by the highly specific genome editing tool CRISPR/Cas9 (D10A) nickase optimized for diatoms (Nawaly et al., 2020). Transformed colonies were respread more than twice on agar plates containing antibiotics to obtain the monoclones. Finally, two lines of independent biallelic knock-out mutants were successfully obtained, denoted as ΔPtBST1_1 and ΔPtBST1_2 (Supplementary Fig. S4A and B). Immunoblotting using the specific PtBST1 antibody indicated that these mutants lacked PtBST1 at the protein level (Supplementary Fig. S4C).

These knock-out mutants, ΔPtBST1_1 and ΔPtBST1_2 showed a trend of slightly retarded growth rate in the LC conditions (Fig. 4A). The growth at the logarithmic phase was evaluated as daily specific doubling rate, the so-called *divisions per day* (Wood et al., 2005), with both mutants showing a significantly lower growth rate (Fig. 4B). Meanwhile, there was no difference in the growth between the wild type (WT) and knock-out mutants in HC conditions (Supplementary Fig. S5A and B).

**Figure 4.**
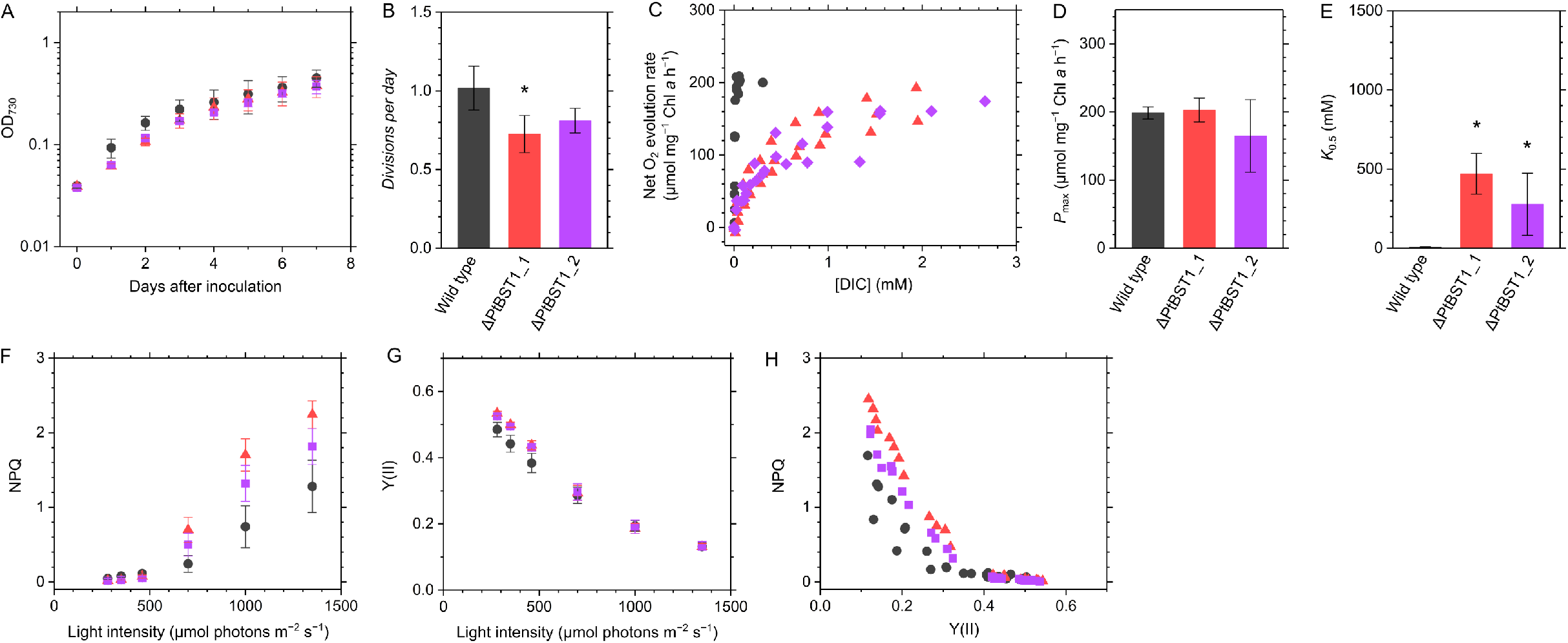
Phenotype of the PtBST1 knock-out mutants grown under air-level CO_2_. (A, B) Growth of *P. tricornutum* wild type (black circles) and the PtBST1 knock-out mutants (ΔPtBST1_1, red triangles; ΔPtBST1_2, purple squares) under air-level CO_2_. Data are shown as the mean ± standard deviation (*n* = 3, biological replicates). Divisions per day was calculated at the logarithmic growth phase. (C−E) Net O_2_ evolution rate at different concentrations of dissolved inorganic carbon (DIC). Cells were illuminated with a white actinic light (800 µmol photons m^−2^ s^−1^). Data of three independent experiments are all plotted. From the kinetics of photosynthetic activity, the maximum O_2_ evolution rate (*P*_max_) and the DIC concentration giving to a half of *P*_max_ (*K*_0.5_) were calculated. (E−G) Chlorophyll fluorescence parameters at different light intensities. Non-photochemical quenching (NPQ) and effective quantum yield of photosystem II, termed as Y(II), were calculated for each sample. Measurements were conducted in the presence of 10 mM NaHCO_3_. Data are shown as the mean ± standard deviation (*n* = 4, biological replicates). (H) Proposed functional model of BST in *P. tricornutum*. Asterisks indicate the significant difference between the wild type and mutants.

### Effects of the lack of PtBST1 on photosynthesis of *P. tricornutum*

We determined the photosynthetic parameters (DIC affinity and maximum rate) by measuring the kinetics of net O_2_ evolution rate over various DIC concentrations, which were quantified with gas-chromatography flame ionization detector (GC-FID). Whereas the maximum photosynthetic O_2_ evolution rate (*P*_max_) was not significantly different between the LC-grown WT and LC-grown mutant cells, at around 200 µmol O_2_ mg^−1^ chlorophyll *a* h^−1^, both LC-grown mutants showed a significantly lower photosynthetic DIC affinities as indicated by the *K*_0.5_ value; that is 0.3−0.5 mM for the knock-out mutants, while in the LC-grown WT cells, it was about 0.01 mM DIC (Fig. 4C−E). These kinetics data clearly indicate that, in BST1-KO mutants, the photosynthetic DIC affinity decreased to 1.5% of that of the WT cells. In the cells grown under HC, there was no difference in either parameter between WT and mutant strains, and both *P*_max_ and *K*_0.5_ were higher than in the cells grown under LC (Supplementary Fig. S5C−E). Importantly, *K*_0.5_ was higher in the HC-grown cells than those in the LC-grown cells even in the absence of PtBST1 (Fig. 4E and Supplementary Fig. S5E).

Non-photochemical quenching (NPQ) was significantly increased to 1.5−2.0 times in both mutants relative to that in the WT cells when the measurement was done under CO_2_ saturating condition (Fig. 4F). This implies that the occurrence of PtBST1 in ST plays a role in suppressing NPQ as long as the CO_2_ supply to the Calvin cycle is not a limiting factor for the linear electron flow of photosystems. Indeed, effective quantum yield value, Y(II) was not significantly different between the WT and mutant cells (Fig. 4G). NPQ and Y(II) inversely correlate during the steady-state photosynthesis under changing actinic light intensity in all strains, but the mutants showed higher NPQ in the inverse relationship (Fig. 4H), strongly suggesting that PtBST1 add some effect to modulate the energy distribution system on the photosystem II. In the HC-grown cells, NPQ was lower in both WT and mutants, compared with the LC-grown cells, and the relationship between NPQ and Y(II) was similar among them (Supplementary Fig. S5F−H).

## Discussion

Aquatic photosynthesis has evolved the biophysical CCM to overcome low CO_2_ availability within the marine environment, with such evolution likely accelerated under the low CO_2_/O_2_ environment after the Carboniferous. Efficient acquisition of abundant external HCO_3−_ (*ca.* 2 mM) results in high accumulations of stromal HCO_3−_, which is eventually converted to CO_2_ by the CO_2_-evolving machinery in the pyrenoid that provides an ample flux of CO_2_ substrate to Rubisco condensate in the pyrenoid (He et al., 2023). In the case of marine diatoms, it is known that the PPT lumen is the site and θ-type CA is the converter in the CO_2_-evolving machinery (Kikutani et al., 2016).

The above-mentioned general model however contains several unsolved problems, the clarification of which is essential to understand the biophysical CCM in diatoms. Of particular importance, i) the mechanism to maintain inner chloroplast structure with the pyrenoid and functionally diverse thylakoid membrane, ii) the system to provide the accumulated stromal HCO_3−_ into the CO_2_-evolving machinery in the PPT lumen, and iii) the mechanism to prevent the efflux of unfixed CO_2_ from the chloroplast, are the immediate issues to be solved.

In the present study, we identified the molecular mechanism responsible for supplying HCO_3−_ to the CO_2_-evolving machinery in the PPT lumen as bestrophin. In a recent study, we found that a complete defect of θ-type CA (Ptθ-CA1) in the PPT lumen in *P. tricornutum* significantly lowered the DIC affinity for photosynthesis (*K*_0.5_ > 2 mM DIC and photosynthesis only saturated at more than 10 mM DIC) (Shimakawa et al., 2023), indicating that the CO_2_- evolving machinery is essential for photosynthesis not only under CO_2_ limited conditions but also even under CO_2_ enrichment 5-times greater than current atmospheric levels. This further implies that the biophysical CCM in marine diatoms is not only necessary under CO_2_ starvation but is required under normal conditions to supply enough CO_2_ to Rubisco to sustain photosynthesis. In support of this, components such as PPT and the luminal Ptθ-CA1 are constitutively expressed regardless of the growth CO_2_ concentrations (Kikutani et al., 2016), while the presence of other components like bestrophinsis clearly stimulated by LC at either transcript or post-transcriptional levels (Fig. 3A and B). This indicates that the diatom biophysical CCM is comprised of two systems; one is the constitutively-operating CCM within the diatom pyrenoid that is an absolutely required to sustain photosynthesis by maintaining, and the other is the LC-inducive CCM that highly strengthens the activity of DIC acquisition and the function of the PPT-based CO_2_-evolving machinery. The latter mechanism is sustained by a few LC inducible CCM factors that in addition to bestrophins include previously identified features such as SLC4 HCO_3−_ transporters at the plasma membrane (Nakajima et al., 2013; Nawaly et al., 2022), and pyrenoidal CAs (Satoh et al., 2001; Harada and Matsuda, 2005; Nawaly et al., 2023).

In diatoms, the mode of operation of their CCM under sub-atmospheric to very low-level CO_2_ (VLC) is totally unknown. In the green alga, *C. reinhardtii*, numerous mutagenesis experiments have indicated that a different type CCM mode operates in LC as compared to VLC. In VLC, *C. reinhardtii* relies more on chloroplast envelope HCO_3−_ transporters, while in LC the CCM primarily utilizes less energy-requiring methods using a CO_2_-leakage barrier, such as LCIB/C complex (putative θ-type CA) from the stroma (Duanmu et al., 2009; Yamano et al., 2010; Fei et al., 2022). Given these examples from green alga, it is very possible that diatoms also operate a further strengthened mode of the CCM under VLC, which would be another interesting topic to be studied. Nonetheless, the phenotype of mutants defective in PtBST1, which showed decreased *K*_0.5_ of LC-grown *P. tricornutum* to approximately 0.5 mM (Fig. 4C and E), clearly indicating that PtBST1 is a component that facilitate the LC-inducible mode CCM but not house-keeping mode of the constitutively-operating CCM in *P. tricornutum* as the photosynthetic DIC affinity of ΔPtBST1 was still much higher than that of the mutants defect of the PPT luminal θ-CA in *P. tricornutum* (Shimakawa et al., 2023).

It should be noted that the system by which BST expression is regulated differs quite markedly between the two species used in the present study. Especially in *T. pseudonana*, it is clear that the production of the *BST* protein is strongly regulated at the translational, rather than transcriptional, level by the ambient CO_2_ concentrations (Fig. 3). CO_2_ responses of diatom CCM factors have been studied mostly at the transcriptional levels (Ohno et al., 2011; Hennon et al., 2015), and thus our data on *T. pseudonana BST* in this study would provide an intriguing new sample to study the post-transcriptional controls of protein expression in response to CO_2_ in diatoms. As for the transcriptional controls, it is known that cAMP plays a role as a secondary messenger to transmit a CO_2_ signal to CO_2_ responsive *Cis*-element via a basic zipper-type transcription factor PtbZIP11 (Harada et al., 2006; Ohno et al., 2011; Hennon et al., 2015) and indeed our present study revealed that the *PtBest1* gene also possess a typical tandem repeat of TGACGT/C motif (Supplementary Fig. S3). This suggests that the *PtBest1* gene is controlled by the same mechanism as *PtCA1* ; the signal control of the secondary messenger cAMP governing the CO_2_/light cross talk (Tanaka et al., 2015). In contrast, the CO_2_-response mechanism to control the expression at the post-transcriptional levels is so far poorly understood in diatoms.

Two-types of subcellular localization were observed among bestrophin isoforms in *P. tricornutum*. Whereas PtBST1 and PtBST2 were localized in the ST outside of the pyrenoid, PtBST3 was specifically localized at/ around PPT (Fig. 2A and B). In *T. pseudonana*, both TpBST1 and TpBST2 were localized at PPT even though a part of TpBST1 also dispersed to the outside of the pyrenoid (Fig. 2C). We initially assumed that such differences of localization reflected the locus-dependent differences in the function of BST factors; that is, BSTs in the PPT would play a specific role in the CO_2_-evolving machinery but ones in the ST membranes would contribute to more general functions of the photosystems such as regulation of thylakoid membrane potential by providing a counter anion to the lumen. However, the striking phenotype of ΔPtBST1 mutants, which totally lacks in the LC-inducible CCM, clearly indicates that even the BST factors outside of the pyrenoid are vital components of the CO_2_-evolving machinery in *P. tricornutum*. In addition, it was also clear that the key enzyme for the CO_2_-evolving machinery, Ptθ-CA1 was very specifically localized at the PPT lumen (Kikutani et al., 2016) and the knockout of Ptθ-CA1 completely abolished not only the LC-inducible CCM but also the constitutive level CCM, clearly indicating the essentiality of the PPT luminal Ptθ-CA1 for the CO_2_-evolving machinery in *P. tricornurtum* (Shimakawa et al., 2023). Given these considerations, we assume that the stromal thylakoid membranes are highly likely to be connected to the PPT in *P. tricornutum* and HCO_3−_ incorporated into the lumen of the ST membranes can readily move to the PPT lumen where it is rapidly dehydrated by Ptθ-CA1 to evolve CO_2_. It is unclear so far whether or not the connection between the ST and PPT is a general feature of diatoms. It is also unclear how the function of PPT-located BST factors differs from those in the ST membranes. It is possible that there are some structural variations regarding the relationship between the ST membranes and PPT membrane among diatoms which might provide slightly different mechanism to the case of *T. pseudonana*. It is also possible that, similar to the dynamics of pyrenoid structure and functional alterations observed in Chlamydomonas, these diatom BST factors might be differentially expressed or localized in response to the severity of CO_2_ starvation, and BSTs in the PPT may function in more severe CO_2_ starvation events. Interestingly, the location of PtBST3 shifted from the PPT to ST by truncating the C-terminal extension of PtBST3 (Fig. 2A). This implies the existence of a unique sequence in the C-terminal extension of PtBST3 targeting the protein to the pyrenoid but we could not find any clear natural disorder sequence nor structural moiety such as amphipathic helix, which was previously reported to occur in pyrenoidal β-CAs (PtCA1 and PtCA2) to be localized in the pyrenoid (Kitao and Matsuda, 2009). A recent study found that BST4 in *C. reinhardtii* has the unique C-terminal region that contains a Rubisco-binding motif and localizes it in the pyrenoid (Adler et al., 2023). Unfortunately, we could not identify the similarity in the C-terminal region between PtBST3 and CrBST4.

The aforementioned second consideration about the functional diversity of BST (i.e., the possibility of BST at the ST membranes as a regulator for the photosystem functions) is still interesting object of this research. Bestrophin comprises a large protein family of anion transporter, which is broadly conserved in bacteria, animals, and plants. It has been reported that several bestrophins can transport HCO_3−_ (Qu and Hartzell, 2008). Transportation of anions across the thylakoid membrane also affects the photosynthetic electron transport, especially at the ST. During the photosynthetic linear electron transport, H^+^ is released in the thylakoid lumen around PSII and at the Q-cycle, storing energy as the proton motive force, which is composed of ΔpH and electric field difference (Δψ) across the thylakoid membranes. In principle, both ΔpH and Δψ are the motive force to drive chloroplast ATP synthase. However, ΔpH is also an important factor in the regulation of light utilization and electron transport in the thylakoid membranes. The luminal acidity activates violaxanthin de-epoxidase and diadinoxanthin de-epoxidase to accumulate zeaxanthin and diatoxanthin, respectively, which is essential to induce dissipation of excess light energy at PSII, the so-called non-photochemical quenching (more specifically qE quenching). Further, the lumen acidification can cause the suppression of electron transport around the cytochrome *b*_6_*f* complex by down-regulating the Q-cycle. To optimize these regulatory mechanisms on the thylakoid membranes, photosynthetic organisms modulate the balance of ΔpH and Δψ. In the C_3_ plant *Arabidopsis thaliana*, the K^+^/H^+^ antiporter KEA3 and the bestrophins (AtBST1 and AtBST2; often referred as another name the voltage- gated Cl^−^ channel VCCN1) function in balancing ΔpH and Δψ (Armbruster et al., 2014; Duan et al., 2016; Herdean et al., 2016). In *P. tricornutum*, the lack of PtBST1 increased NPQ (Fig. 4F), probably due to the decreased consumption of H^+^ in the PPT lumen for the dehydration of HCO_3−_ (Kikutani et al., 2016; Shimakawa et al., 2023). Alternatively, spontaneous dehydration of lumen-imported HCO_3−_ in the ST membranes simply accelerated by luminal acidity for some extent even in the absence of CA, which may have moderate effect on suppressing ΔpH. Either or both these mechanisms could function in modulating the extent of NPQ depending on the CO_2_ availability.

In the present study, the most abundantly expressed PtBST1 was shown to be bifunctional factor, which sustains the LC-induced level of the CCM in *P. tricornutum*, most probably as a part of the CO_2_-evolving machinery of the pyrenoid, and also modulates the ΔpH in the entire thylakoidal photoreaction. The study also discovered the diverse localization of multiple LC-inducible BST factors in the chloroplast of both *P. tricornutum* and *T. pseudonana*. The future study on the function and regulation of these BST factors in response to the different level of CO_2_ starvation would provide knowledge on dynamic change in the mode of the CO_2_- evolving machinery in the pyrenoid-based CCM in diatoms.

## Methods

### Cultures

The marine diatoms *P. tricornutum* Bohlin (UTEX642) and *T. pseudonana* (Hustedt) Hasle et Heimdal (CCMP 1335) were axenically and photoautotrophically cultured in artificial seawater medium with the addition of 0.31% half-strength Guillard’s ‘F’ solution (Guillard and Ryther, 1962; Guillard, 1975) supplemented with 10 nM sodium selenite under continuous light (20°C, 40 μmol photons m^−2^ s^−1^, fluorescent lamp). The cultures were aerated with ambient air (0.04% CO_2_) or 1% CO_2_ gas. For the culture of *T. pseudonana*, the concentration of NaCl was lowered to 270 mM in the medium. The growth of diatoms was evaluated as the increase in optical density at 730 nm.

### Phylogenetic analysis

Amino-acid sequences of bestrophin family proteins were collected from DiatOmicBase (https://www.diatomicsbase.bio.ens.psl.eu/), JGI Genome Portal (https://genome.jgi.doe.gov/portal/), and NCBI (https://www.ncbi.nlm.nih.gov/). For PtBST1, PtBST2, PtBST4, and TpBST1, the correct full-length sequences were determined for the laboratory strains using a SMARTer RACE 5’/3’ kit (Takara, Shiga, Japan). These sequences were aligned by ClustalW, and a conserved domain was defined in reference to the motif of PtBST1. To this motif, the evolutionary history was inferred by using the Maximum Likelihood method and Le_Gascuel_2008 model (Le and Gascuel, 2008). Initial tree(s) for the heuristic search were obtained automatically by applying Neighbor-Join and BioNJ algorithms to a matrix of pairwise distances estimated using the JTT model, and then selecting the topology with superior log likelihood value. A discrete Gamma distribution was used to model evolutionary rate differences among sites (5 categories [+G, parameter = 1.6226]). All positions containing gaps and missing data in the amino-acid sequence alignment were eliminated. Evolutionary analyses were conducted in MEGA X (Kumar et al., 2018).

### Expression of GFP fusion proteins

The full-length coding regions of PtBST1, PtBST2, PtBST3, PtBST4, TpBST1, and TpBST2 were amplified by PCR with the primers shown in Supplementary Table S2. The fragments of PtBST1 and PtBST2 were cloned into the *Eco*RI restriction site in pPha-T1/*egfp* vector (Tachibana et al., 2011) using Mighty Mix (Takara). PtBST3 and PtBST4 fragments were cloned using Gibson assembly system (New England Biolabs, Ipswich, MA, USA) into the *Eco*RI restriction site in pPha-NR/*egfp* vector that was developed by replacing *fcpA* promoter in pPha-T1/*egfp* vector with nitrate reductase promoter. The resulting plasmid for PtBST3:GFP was linearized by inverse PCR with the primers shown in Supplementary Table S2, and then ligated with Mighty Mix to generate the plasmid for PtBST3^Δ407−519^:GFP. The coding regions of TpBST1 and TpBST2 were cloned with Mighty Mix into the *Eco*RV restriction site in pTpNR/*egfp* vector (Poulsen et al., 2006).

The resulting plasmids were introduced into the *P. tricornutum* or *T. pseudonana* cells by biolistic particle bombardment (PDS-1000/He, Bio-Rad, Hercules, CA, USA) as previously described (Zaslavskaia et al., 2000). Transformants were screened on 1.2% (w/v) agar medium containing 100 μg mL^−1^ zeocin (Invitrogen, Waltham, MA, USA) or 100 μg mL^−1^ nourseothricin (Jena Bioscience, Germany).

### Confocal fluorescence microscopy

The transformant cells grown in liquid medium were collected at the logarithmic growth phase for the confocal fluorescent microscopy with TCS SP8 (Leica, Wetzlar, Germany). Chlorophyll autofluorescence was evaluated from the emission between 600 and 750 nm excited by a 552 nm laser. GFP was excited by a 488 nm laser, and the green fluorescence was detected at 500−520 nm (Shimakawa et al., 2022).

### Immunoelectron microscopy

The transformant of *P. tricornutum* that expresses PtBST1 tagged to GFP was fixed by the procedure previously described (Kikutani et al., 2016). Thin sections were cut with an RMC MT-X ultramicrotome (RCM, Tucson, AZ, USA) and mounted on nickel slot grids, followed by an edging step with 1% (w/v) hydrogen peroxide. After the blocking step, the sections were reacted with polyclonal anti-GFP antibody (AnaSpec, Fremont, CA, US) diluted 1:500 in 3% (w/v) BSA in PBS at 25°C overnight. After rinsing with PBS, they were incubated for 60 min at room temperature with a goat anti-rabbit IgG conjugated to 10-nm colloidal gold particles (1:50 diluted in PBS; BBI Solutions, Crumlin, UK). The thin sections were stained with TI blue (Nisshin EM, Aichi, Japan), following washing with distilled water. The sections were observed with a JEM-1011 electron microscope (JEOL, Tokyo, Japan).

### qRT-PCR

Cells grown under 0.04% and 1% CO_2_ were harvested and frozen in liquid nitrogen at the logarithmic growth phase, which were kept at −80°C until total RNA and protein extractions as described below. Total RNA was extracted from the frozen cells to prepare cDNA following the method previously reported (Samukawa et al., 2014). qRT-PCR was performed with GeneAce SYBR qPCR Mix α No ROX (Nippon Gene, Tokyo, Japan) in Thermal Cycler Dice Real Time System II (Takara). Each *Actin1* genes (*PtActin1* and *TpActin1*) were used as the reference genes. All primers used were shown in Supplementary Table S3.

### Immunoblotting

The cells were disrupted by a UD-201 sonicator (duty, 40; output 0.5; TOMY, Tokyo, Japan) in 50 mM Tris-HCl (pH 6.8) containing 10% (v/v) glycerol and 1% (w/v) SDS for 2 min on ice. Unbroken cells were removed by centrifugation at 2,000 × *g* for 5 min, and the resulting supernatant was analyzed by sodium dodecyl sulfate polyacrylamide gel electrophoresis (SDS-PAGE). Following electrophoresis, the proteins were blotted onto a polyvinylidene fluoride membrane and subsequently labelled with custom polyclonal antibodies (Japan Bio Serum, Hiroshima, Japan) specific to the target sites of PtBST1 (GDTTGITMDQPHNA), TpBST1 (AGQGQQEYTEENA), and TpBST2 (WRGQGLDKEEQQY). After washing with a phosphate buffer saline containing 0.05% (v/v) Tween 20, bound antibodies were revealed with a peroxidase-linked secondary anti-rabbit antibody (Promega, Madison, WI, USA) and visualized by chemiluminescence (ImmunoStar Zeta, Wako, Osaka, Japan). RbcS was detected as a control with a rabbit anti-RbcS antiserum against the 35−50 region of the amino acid sequence in both *P. tricornutum* and *T. pseudonana* (diluted 1:2000; Sigma, St. Louis, MO, USA).

### Genome editing of *PtBST1*

The nucleotide locations 1014–1033 and 1045–1064 in *PtBST1* (*Phatr3_J46336*) were chosen as the target pair of single-guide RNA. The PCR fragment including them were prepared with pPt_dual_sgRNAs as the template according to the method previously reported (Nawaly et al., 2020), and then ligated into the BsaI site of pAC-ctCRISPR-Cas9n-2 (Shimakawa et al., 2023). The resulting plasmid vector was introduced into *P. tricornutum* WT cells by bacterial conjugation with *E. coli* S17-1. Transformants were screened on agar plates supplemented 100 μg mL^−1^ Zeocin (Invitrogen, Waltham, MA, USA), and the candidate colonies, which were selected with fragment size assay of target sequence by genomic PCR, were re-spread twice under HC condition. Direct sequencing was performed to confirm whether or not each colony were monoclone.

### O_2_ measurement

Wild type and the mutant cells of *P. tricornutum* were harvested at the logarithmic growth phase and resuspended in a DIC-free F/2 artificial water freshly prepared. Chlorophyll *a* concentration of the samples was determined in 100% (v/v) methanol (Jeffrey and Haxo, 1968), and the cell samples were applied to oxygen electrode (Hansatech, King’s Lynn, U.K.) at 10 µg chlorophyll *a* mL^−1^ in the DIC-free artificial sea water (pH 8.1). Simultaneous measurement of net O_2_ evolution rate with total DIC concentration in the sample mixture was achieved by the combination of oxygen electrode measurement with GC-FID analysis (GC-8A, Shimadzu, Kyoto, Japan) during a stepwise addition of NaHCO_3_ as previously reported (Kikutani et al., 2016). The photosynthetic parameters were calculated from the plot of O_2_ evolution rate against DIC concentration by curve fitting with the non-linear least squares method: *P*_max_, maximum net O_2_ evolution rate; and *K*_0.5_, DIC concentration giving a half of *P*_max_.

### Chlorophyll fluorescence analysis

Pulse-modulated excitation was achieved using an LED lamp with a peak emission of 625 nm in a Multi-Color-PAM (Walz, Effeltrich, Germany). Pulse-modulated fluorescence was detected within the range of wavelength limited by RG 665 long pass and SP 710 short pass filters. Cells were illuminated with white actinic light from the LED array at 20°C. The effective quantum yield of PSII, Y(II), was calculated as (F_m_ʹ – Fʹ)/F_m_ʹ with: F_m_ʹ, maximum fluorescence from light-acclimated cells; Fʹ, fluorescence emission from light-acclimated cells. Non- photochemical quenching (NPQ) was calculated as (F_m_ – F_m_ʹ)/F_m_ʹ with: F_m_, maximum fluorescence from dark-acclimated cells. Short saturation flashes (10,000 µmol photons m^−2^ s^−1^, 600 ms) were applied to determine F_m_ and F_m_ʹ.

## Author contributions

Y.M. conceived the research plan; M.N. and G.S. performed phylogenetic analysis; M.N., K.Y., R.A., and S.I. analyzed subcellular localizations of GFP-fused proteins with the assistance from G.S. and Y.T.; M.N., K.Y., and R.A. analyzed the gene and protein expressions; M.N. generated the knock-out mutant and performed photosynthetic measurements with the assistance from G.S.; C.N. performed immunoelectron microscopy; G.S. and Y.M. mainly wrote the manuscript.

## Acknowledgements

We wish to thank Mrs. Nobuko Higashiuchi and Mrs. Eri Nakayama (Kwansei-Gakuin University) for their technical assistance, and Dr. Matthew B. Brown (Kwansei-Gakuin University) for kindly proofreading our English writing. Further, we thank Dr. Luke C. M. Mackinder and Dr. Onyou Nam (University of York) for their critical reading and helpful comments.

## Supplementary materials

**Supplementary Fig. S1.**
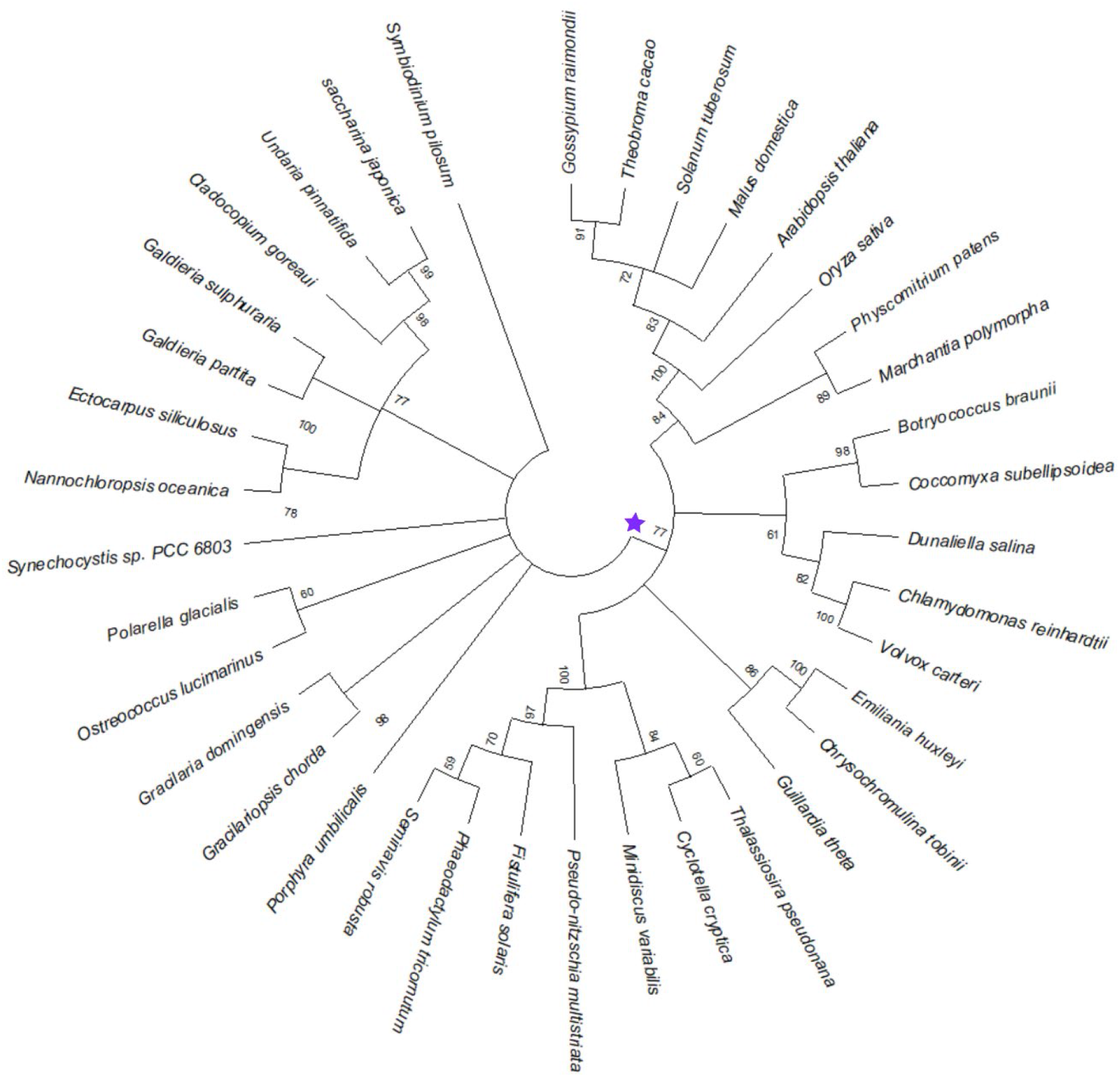
Phylogenetic tree of bestrophin family proteins in photosynthetic organisms. The image was presented as a bootstrap consensus tree in MEGA X. Branches corresponding to partitions reproduced in less than 50% bootstrap replicates are collapsed. One each BST ortholog per species were selected for the phylogenetic analysis, based on the similarity to PtBST1. Homologs in the BST clade (indicated by purple star) was further analyzed (see the main text).

**Supplementary Fig. S2.**
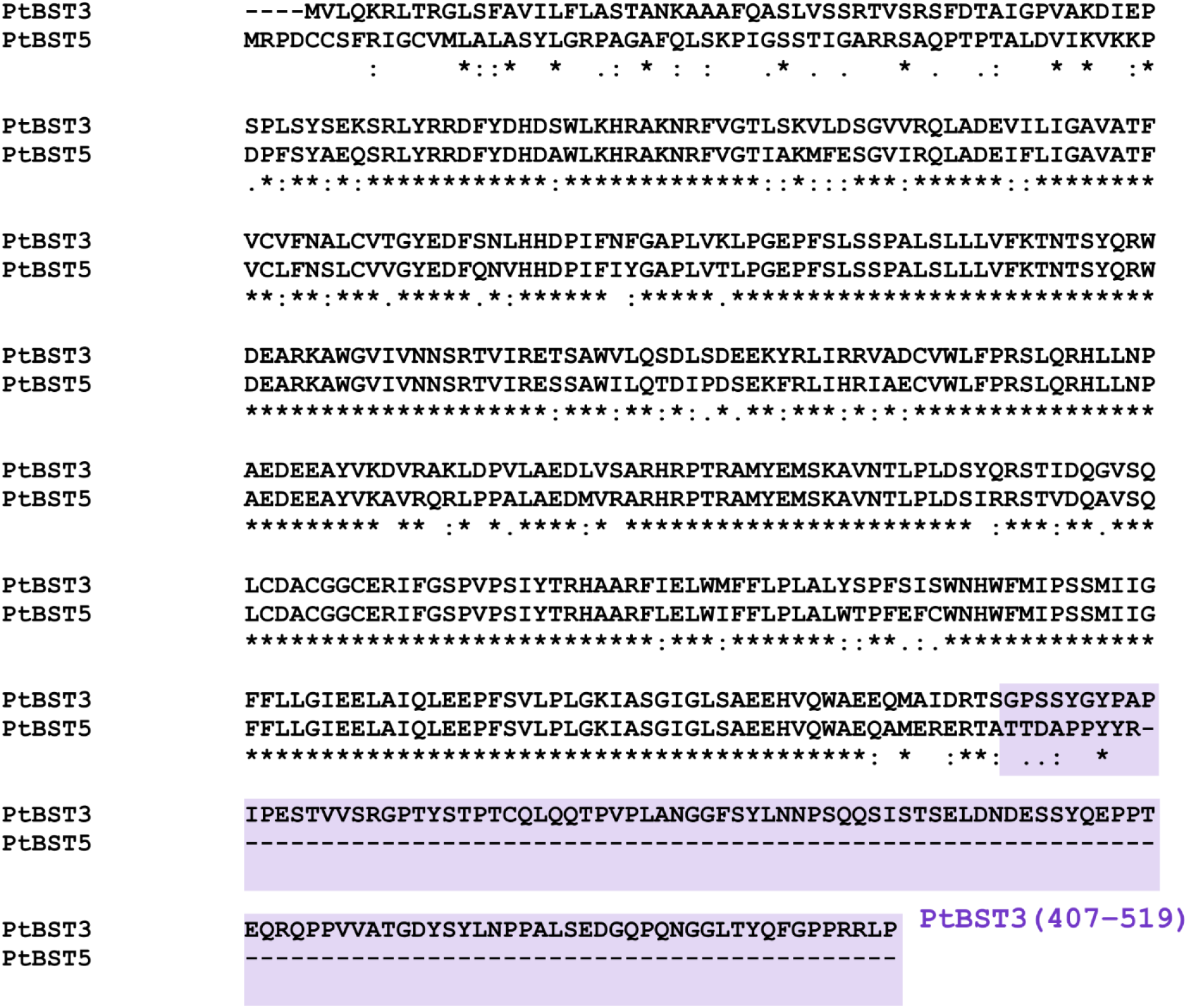
Amino-acid sequence alignment between PtBST3 and PtBST5 by ClustalW. The C-terminal region of PtBST3 (407−519) is shown in the purple shading.

**Supplementary Fig. S3.**
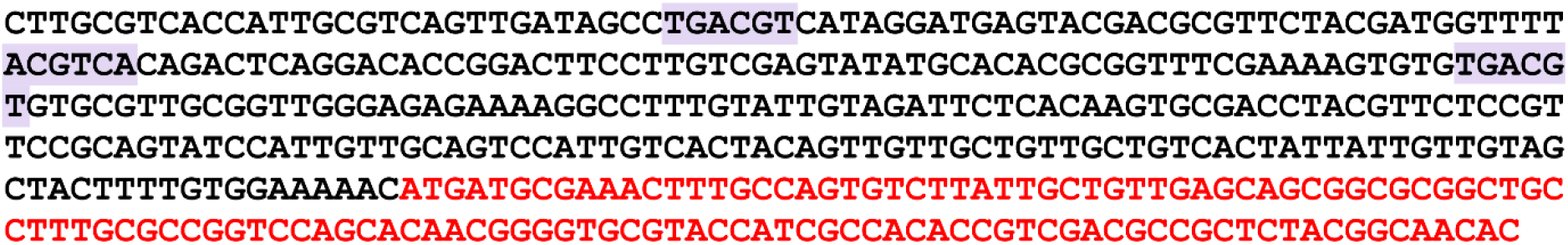
Genome DNA sequence of the PtBST1 upstream region in *P. tricornutum*. Three CO_2_-cAMP-responsive elements (CCRE) were identified as indicated by purple shadings. The coding sequence of PtBST1 is highlighted in red.

**Supplementary Fig. S4.**
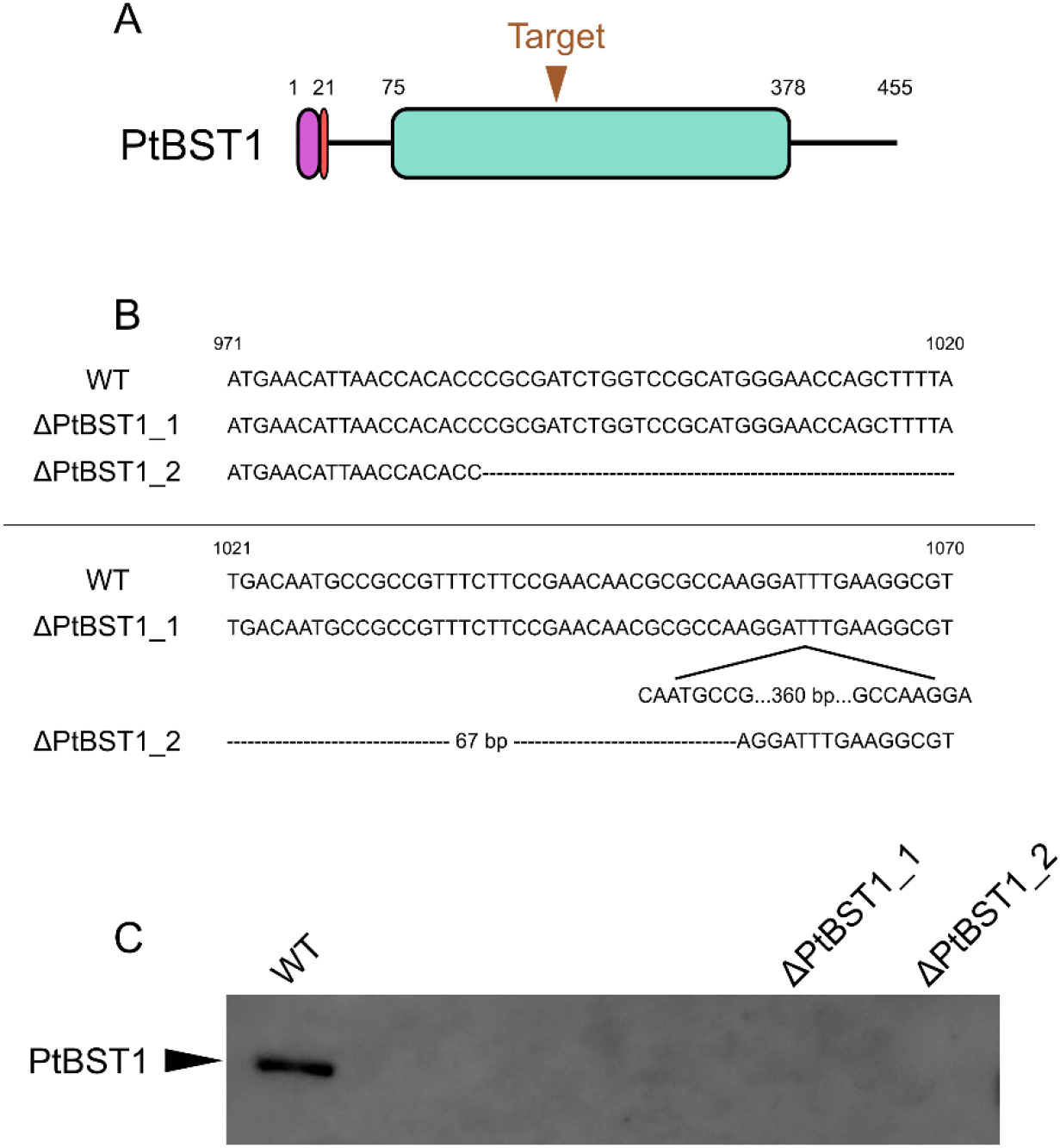
Genome editing by CRISPR Cas9 nickase to PtBST1 in *P. tricornutum*.(A) Target site of gRNAs in PtBST1. Blue box shows the conserved motif in bestrophins. (B) DNA sequence alignment of *PtBST1* in WT and each knock-out mutant. (C) Western blot analysis of WT and knock-out mutants using anti-PtBST1 antibody.

**Supplementary Fig. S5.**
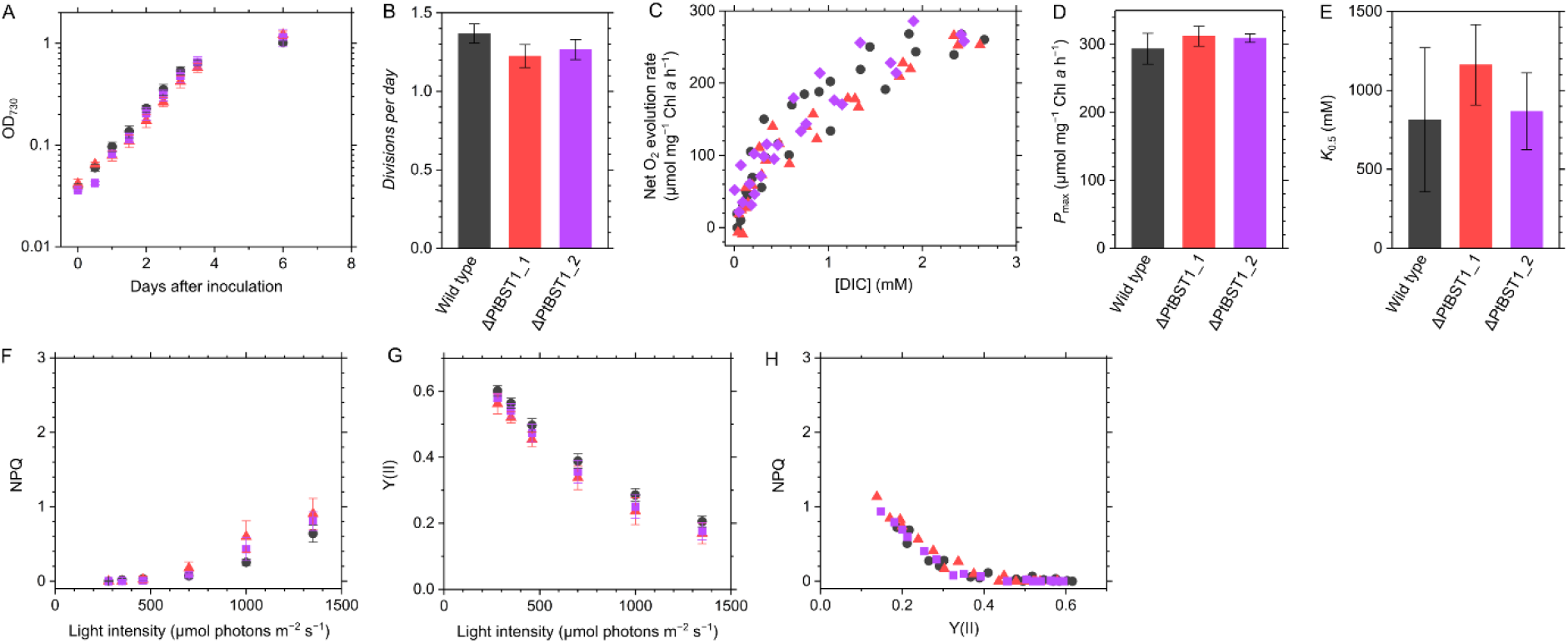
Phenotype of the PtBST1 knock-out mutants grown under high CO_2_. (A, B) Growth of *P. tricornutum* wild type (black circles) and the PtBST1 knock-out mutants (ΔPtBST1_1, red triangles; ΔPtBST1_2, purple squares) under high CO_2_. Data are shown as the mean ± standard deviation (*n* = 3, biological replicates). Divisions per day was calculated at the logarithmic growth phase. (C−E) Net O_2_ evolution rate at different concentrations of dissolved inorganic carbon (DIC). Cells were illuminated with a white actinic light (800 µmol photons m^−2^ s^−1^). Data of three independent experiments are all plotted. From the kinetics of photosynthetic activity, the maximum O_2_ evolution rate (*P*_max_) and the DIC concentration giving to a half of *P*_max_ (*K*_0.5_) were calculated. (F−H) Chlorophyll fluorescence parameters at different light intensities. Non-photochemical quenching (NPQ) and effective quantum yield of photosystem II, termed as Y(II), were calculated for each sample. Measurements were conducted in the presence of 10 mM NaHCO_3_. Data are shown as the mean ± standard deviation (*n* = 4, biological replicates).

**Supplementary Table S1.** Chloroplast transit peptides predicted by SignalP in BST isoforms in *P. tricornutum* and *T. pseudonana*.

| Name | Protein ID | ER-SP<sup>a</sup> | ASA-FAP<sup>b</sup> |
| --- | --- | --- | --- |
| PtBST1 | 46336 | + | AAA-FAP |
| PtBST2 | 26635 | + | TQA-FIV |
| PtBST3 | 46366 | + | AAA-FQA |
| PtBST4 | 46360 | + | ADA-FQA |
| TpBST1 | 4819 | + | TSA-FST |
| TpBST2 | 4820 | + | ISA-FAP |

The sequence data were obtained at DiatOmicBase (https://www.diatomicsbase.bio.ens.psl.eu/) or JGI Portal (https://genome.jgi.doe.gov/portal/), or experimentally determined by 5'-RACE. <sup>a</sup>The presence of endoplasmic reticulum signal peptide (ER-SP) was predicted by SignalP-6.0 (https://services.healthtech.dtu.dk/services/SignalP-6.0/). <sup>b</sup>The plausible ASA-FAP motif for targeting to chloroplasts was manually defined in the region just after the ER-SP site.

**Supplementary Table S2.**
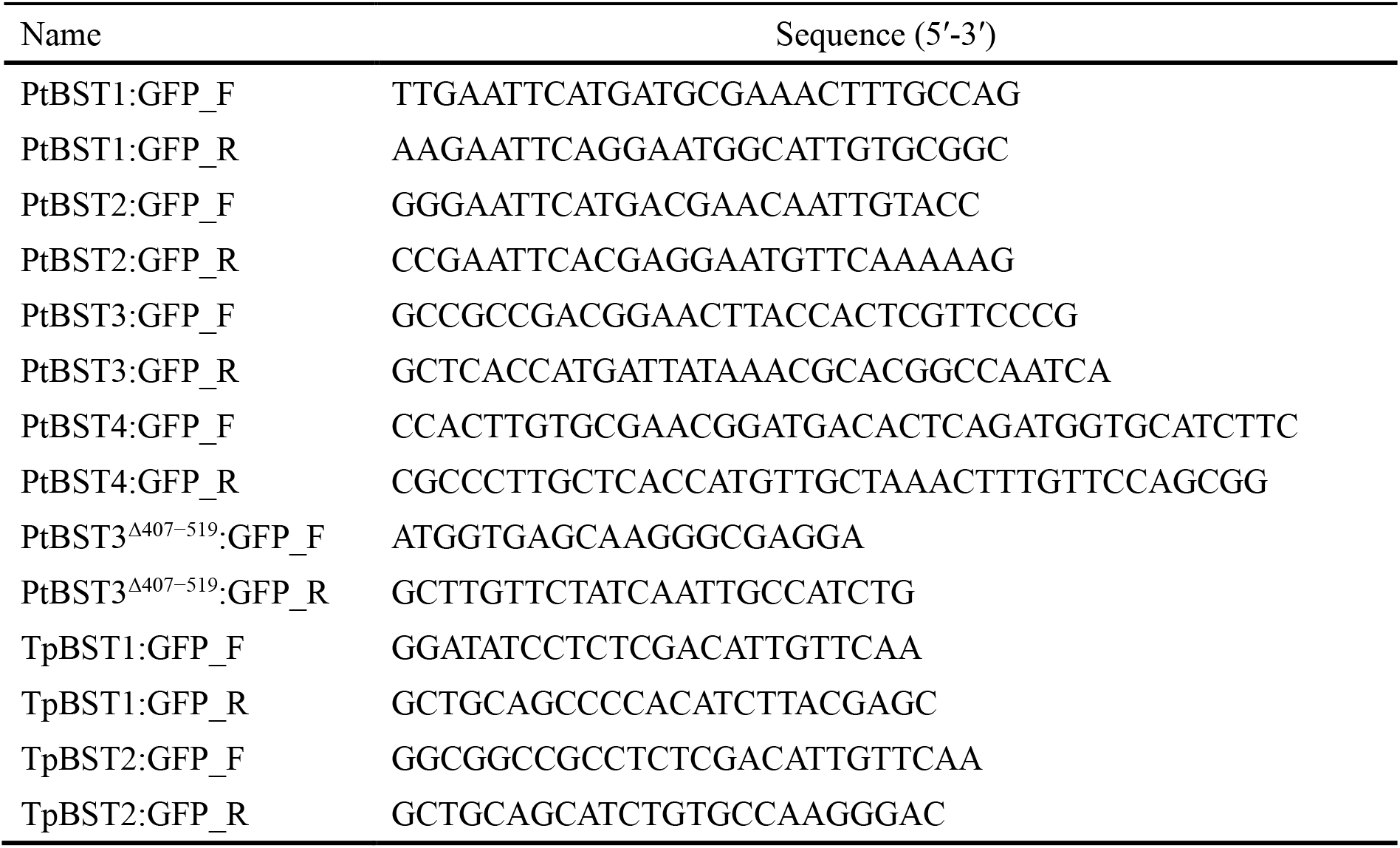
Primers used for the plasmid construction for the expression of GFP-fused proteins.

**Supplementary Table S3.**
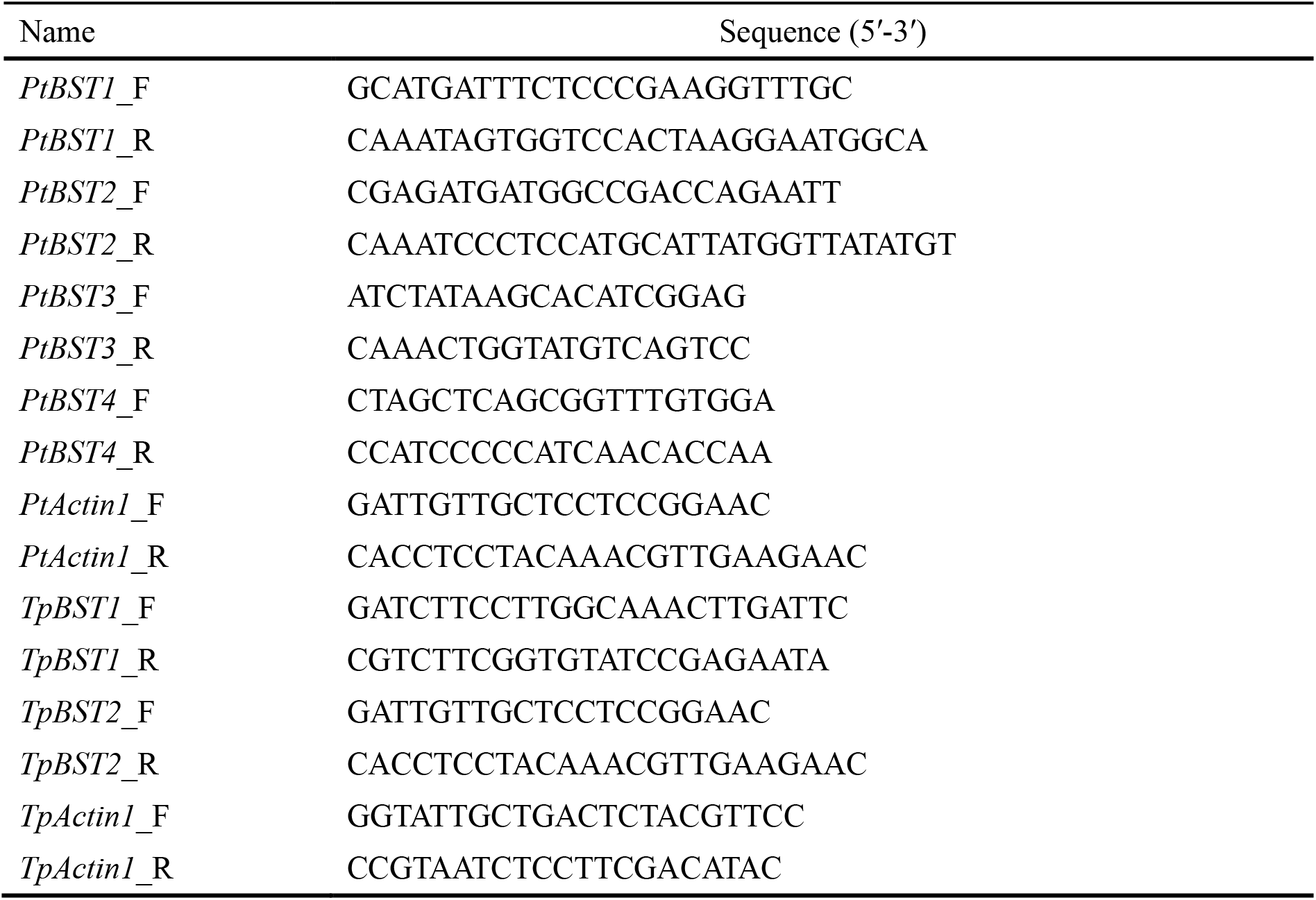
Primers used for qRT-PCR.

**Supplementary Table S4.**
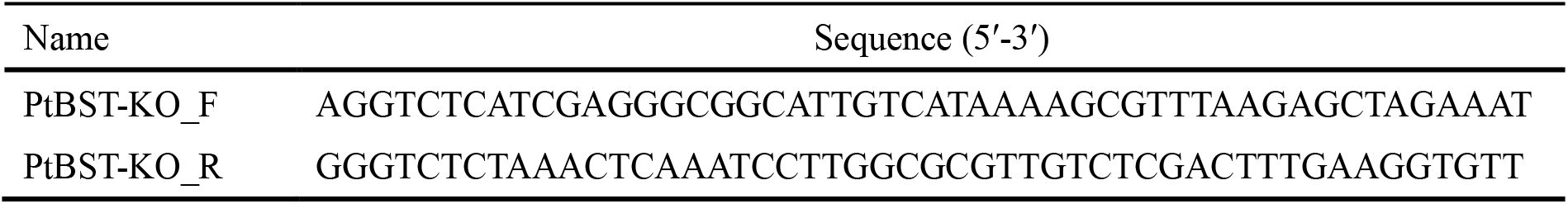
Primers used for the plasmid construction for genome editing by CRISPR-Cas9 nickase.

